# Cryopreservation Enables Long-Term Studies of *Octopus vulgaris* Hemocyte Immune Functions

**DOI:** 10.1101/2024.07.19.604316

**Authors:** MM Costa, E Paredes, M Peleteiro, F Gambón, S Dios, C Gestal

## Abstract

As with their nervous system and other physiological traits, the immune system of cephalopods, in general, and of the common octopus, Octopus vulgaris, in particular, could also present highly evolved characteristics compared to other classes of molluscs. However, to date, there is not much information about it, and studying the defense mechanisms is a key step in understanding their response system to external aggressions and, thus, having the tools to anticipate animal health problems and ensure their welfare.

The lack of cell cultures in molluscs is a major problem when carrying out *in vitro* assays that help to deepen the knowledge of the main immune cells of this species. Cryopreservation becomes an alternative to maintain viable and functional cells after freezing/thawing processes, allowing a larger repertoire of studies to be performed with the same sample or to perform time courses.

Having access to good quality cells for long periods will allow to cover a larger repertoire of studies with the same sample or to perform time courses, avoiding logistic bottlenecks that could arise due to the loss of viability and/or functionality of the cells on the lab bench, the lack of time to develop all the desired assays in the same day or the possible losses derived from transporting the samples between different laboratories. Additionally, high-quality suspensions of viable and functional cells are required for successful massive sequencing studies, such as single-cell analysis, where these aspects are the key to an optimal identification.

We show here the first functional results from cryopreserved octopus hemocytes, where the cells were able to maintain viability above 80% after two months post cryopreservation and storage at −80°C and their functional ability to phagocytose bacteria similar to fresh cells. Our data revealed that the acclimation process after thawing was essential to recovering the functional activity of the cells.

The results presented here will facilitate the study of the functions of the most important immune cells of this species and will provide tools for cell preservation in other molluscs species.

## INTRODUCTION

The global market of cephalopods is an important business that generates multi-billion dollars annually and contributes to the livelihoods of thousands of coastal communities around the world. Despite a global trend of decreasing cephalopod has been observed since 2014, their commercialization in terms of volume (tonnes) and economic yield (USD) has shown a consistent upward trend since the 1950s. The main countries involved in cephalopod extraction are China, India, Argentina, and Peru. The primary import markets for cephalopods include Asian countries (China, Japan, Thailand, Republic of Korea, Vietnam, and India), as well as Italy, Spain (within the European Union), Morocco, and the USA (FAO, 2024; Ospina-Alvarez et al., 2022). The commercialization of octopus is indeed increasing in the USA. Recent reports indicate a growing demand for octopus in this country, driven by the popularity of ready-to-eat and high-quality seafood products (USD Analytics, 2024). Specifically, the common octopus (*Octopus vulgaris*) is a highly sought-after species, which is marketed as a fresh, frozen and chilled product, due to the high quality of its meat and Spain is one of its main consumers (g/capita/day). This fact makes Spain one of the most important markets of this species and the main European trader to meet the growing worldwide demand (Ospina-Alvarez et al., 2022). This high demand, together with its short life cycle, high growth rate, easy adaptation to captive conditions, high protein content and high market value, make octopus an ideal candidate for aquaculture diversification (Powell, 2022), with a cycle that has already been closed in captivity (Tur et al., 2020) and is being transferred to an industrial scale.

Cephalopods in general, and common octopus in particular, have been considered an enigmatic group of animals that have evolved until acquiring very high capacities. Despite belonging to the invertebrate group, cephalopods represent the most advanced class of the phylum Mollusca showing innovative traits (Tanner et al., 2017). They are phylogenetically placed in the most evolved branch of molluscs and, specifically, the common octopus belongs to the family of coleoids, which lack the typical shell of molluscs (Whalen and Landman, 2022). This loss of the shell has made octopuses acquire characteristics that differentiate them from other molluscs and that allows them a life much more evolved life to thrive in the environment in which they live. Such is the case with their movements of propulsion, their camouflage through special cells (chromatophores) or their high vision capacity, the efficiency of the movement of their arms, their highly developed nervous system with learning capacity, or the presence of a wide neuronal network with a very complex brain and a centralized nervous system (Hochner, 2008; Gutnick et al., 2020; Shook et al., 2024). The presence of all these highly evolved features concerning other species of the same phylum made it in an excellent model species for multiple physiological and neurological biomedical studies (Di Cosmo et al., 2018), as well as for behavioral research (Grant et al., 2006; Hochner, 2008).

As with their brain, nervous system, and other physiological components, the immune system of these organisms, despite being an innate immune system as invertebrates, may present more advanced characteristics than those of other related species. However, the knowledge of the immune system of cephalopods is nowadays limited. Further studies and the identification of the molecular and functional mechanisms that regulate their defense mechanisms will allow to advance into the knowledge of this species. The privileged phylogenetic situation of this species (the most evolved group of invertebrates) also makes it an excellent model for comparative immunology studies. The application of massive sequencing technologies has offered novel very valuable information suggesting that a complex immune response might be operating in these animals (Castellanos-Martinez et al., 2014a; Gestal and Castellanos-Martínez, 2015; Castillo et al., 2015; Albertin et al., 2015; Maselli et al., 2020; Pérez-Polo et al., 2023). However, further studies to deep into the knowledge of their immune system are needed.

Knowing the mechanisms by which this species immune system recognizes foreign organisms and triggers a response against them, becomes one of the keypoints to understand how its defense system works. It is important to note that the octopus, as an invertebrate animal, does not possess an adaptive immune system, capable of creating a secondary response based on the production of antibodies and adaptive memory, but simply possesses an innate system that constitutes its first line of defense against pathogens and microorganisms. This system, although more primitive at the phylogenetic level, has highly efficient strategies, from the simplest mechanisms such as the production of toxic oxygen or nitrogen radicals (Castellanos-Martinez et al., 2014b), to the recognition of pathogen-associated molecular patterns (Castellanos-Martinez et al., 2014a) and the triggering of cell signaling pathways that ultimately activate the expression of specific effector genes (Castellanos-Martinez et al., 2014a; Gestal and Castellanos-Martínez, 2015; Castillo et al., 2015; Albertin et al., 2015; Pérez-Polo et al., 2023). Although these mechanisms have already been described in the common octopus and several effector and modulatory genes have been reported, it is necessary to carry out in-depth studies to understand each cell type (both at the molecular and functional level) and be able to explore these functions in detail. In this sense, single-cell RNA sequencing (scRNA-seq) analysis becomes a powerful tool that allows the study of gene expression at the single-cell level. Unlike traditional RNA sequencing methods that analyze bulk populations of cells, scRNA-seq provides much higher resolution, revealing cellular heterogeneity within a sample (Lähnemann, et al., 2020). However, to successfully perform scRNA-seq analysis with reproducible and high-quality results, it is crucial that the starting cell population strictly meets several requirements such as; a high cellular viability (as dead or damaged cells can release degraded RNA that affects data quality), a high number of cells is needed to ensure adequate and representative coverage of the cell population, a high quality of the cell population where the cells are completely dissociated, without the presence of aggregates and in a free of contaminants and cellular debris medium that respects their integrity. Moreover, cells must be carefully handled during all steps of the process to ensure their functional capabilities and minimize cell death and stress, which could potentially alter gene expression profiles, so the choice of medium becomes another fundamental requirement of this process.

Consequently, having an inexhaustible source of biological material from cell cultures could be a great tool. Not only to facilitate laboratory tasks and increase biological replication, but also as a replacement technique to contribute to the achievement of the 3Rs goal in animal experimentation. However, one of the main problems in carrying out functional and molecular studies in molluscs cells is the absence of immortal cell lines. Although primary cultures have been achieved in some species up to 40 days old (Troncone et al., 2014; Ji et al., 2017; Ladhar-Chaabouni et al., 2021) there is no hemocyte cell line that would allow less limited studies to be carried out, where uniformity between replicates, adequate sample size or continuous availability of replicates would greatly facilitate the experimental design of many assays. Furthermore, it must be taken into account that, in addition to being cells that cannot withstand long periods in primary cultures, one of the defense strategies of these invertebrate organisms is based on the capacity of aggregation of their main phagocytic cells: the hemocytes, which, in many cases, makes it impossible to carry out long-term assays. Moreover, for many techniques, the presence of individual cells is recommended (e.g., flow cytometry) or, such as it has been mentioned above, even mandatory (e.g., sc-RNA seq). Keeping cells in primary culture increases the chances of these aggregations (even when using anti-aggregation solutions) and makes it difficult to carry out this type of technique without interferences. Consequently, in the absence of cell cultures that keep cells viable and individualized for long periods, it is necessary to design other strategies that allow the analysis of a wide range of cell parameters in a feasible way both in time and logistics.

Cryopreservation, therefore, becomes an excellent alternative for freezing and thawing cells while maintaining their integrity and functionality. Although several invertebrate marine species have already standardized their cryopreservation protocols for sperm cells or even for whole embryos and larvae (Vacquier and Hamdoun; 2024; Lago and Paredes, 2023; Anjos et al., 2022), a unified protocol is not available for different cell or tissue types, but rather each step of the process needs to be optimized, from the choice of cryoprotectant agent (CPA) and study of its toxicity effects to the freezing/thawing conditions. The high diversity of marine species and their differences in form, complexity, size, or even in living/feeding/reproducing mechanisms, make this process extremely meticulous, requiring detailed studies of each stage, with a high risk of loss of cell viability if any step of the process is not fully optimized. Even in cells from the same organism, conditions can vary radically and the same CPA or freezing/thawing method can be toxic or lethal for some cells and not for others (Campos et al., 2024; Olver et al., 2023). For instance, in the field of marine biology, there are multiple studies on sperm cryopreservation in different species, while the process of preserving oocytes has not yet been optimized.

The applications of the use, study, and culture of marine invertebrates are wide. Different utilities have been described from the biomedical, physiological, ecological, toxicological, or socioeconomic point of view (Wolkers and Oldenhof; 2015). Therefore, the applications of cryopreservation are as diverse as the applications of each of the cryopreserved species. Specifically, in the field of aquaculture, cryopreservation has already contributed to the implementation of selective breeding programs, to the preservation of biodiversity through biobanks, and to improve the hatchery spat overcoming the limitations of this important industry (Paredes et al., 2022; Heres et al., 2022). Despite the wide range of applications offered by this discipline, to our knowledge, the cryopreservation protocol of any immune cell in marine invertebrates species has not yet been optimized, making it difficult to study their identification, both molecular and functional, and keeping closed a door that would allow many studies that, to date, have not been possible.

In the present work, we present the first results of long-term storage of octopus hemocytes. To our knowledge, this is the first work in which a marine invertebrate immune cell is successfully cryopreserved. These results open up a wide range of opportunities for the study of these key cells in the defense system of these organisms. Having a source of live and functional cells makes it possible to carry out experimental assays that would be impossible to perform in a single day, increasing the number of variables or sampling times to be performed, or sending cells for analysis between laboratories. Taking into account the high degree of specialization of these organisms and the place that this species occupies in evolution, being able to maintain cryopreserved cells for long periods of time will help to have an experimental model that can be used not only to study the welfare and health of these animals, but also from a biomedical and applied point of view. Identifying molecules with bactericidal, antiviral or antitumor activity are some of the challenges that could be studied with cryopreserved cells. It is also important to note that the ability to preserve these cells in an active functional state will also help lay the groundwork for the development of a possible future cell line that would be an important step in the methodology of invertebrate’s immunology.

In our opinion, this tool will provide advantages for the study of the most important cells involved in the defense against pathogens in *Octopus vulgaris*, facilitating the knowledge of the defense mechanisms of this species with enormous potential as model species and also for aquaculture diversification, which will help to implement improvements in their culture conditions and to evaluate possible strategies to preserve its health and welfare.

## 2. MATERIAL AND METHODS

### 2.1. Animal sampling and care

O. vulgaris specimens (measuring 1.5 kg on average) were collected by traps, an artisanal fishing gear used by local fishermen from the Ria of Vigo, Spain (24° 14.09′N, 8° 47.18′W). Animals were properly transported to the Experimental Culture Facilities of IIM-CSIC (center for the breeding and use of experimental animals under the REGA code ES360570202001). Transport, housing, handling, and experimentation were performed according to the Spanish law RD53/2013 within the framework of the European Union directive on animal welfare (Directive 2010/63/EU) for the protection of animals employed for experimentation and other scientific purposes, following the Guidelines for the care and welfare of cephalopods published by Fiorito et al. (2015), and approved by the Ethic Committee of the National Competent Authority (CEIBA2014-0108; ES360570202001/24/EDUCFORM 07/CGM01). Individuals were maintained in tanks of 500 L of filtered aerated seawater at 15 ± 1°C with a continuous re-circulating flow. The photoperiod was 12 h light:12 h dark and cleaning and parameter checks were performed every day. They were fed daily with frozen fish and fresh mussels. Before the experimental trial, octopuses were acclimatized for one week. All animals were individually stabled due to their territorial and cannibal nature.

### 2.2. Hemolymph collection

Animals were bath anesthetized by immersion in 3L of a 1.5% MgCl_2_ solution dissolved in filtered seawater (FSW) supplemented with 1% of ethanol (Fiorito et al., 2015). Hemolymph samples were collected from the caudal vein using a disposable syringe previously semi-filled with marine antiaggregant solution (MAS) (Troncone et al., 2014) to dilute the hemolymph in a proportion 1:1 and avoid the cellular aggregation of the collected hemocytes. Samples were immediately centrifuged at low speed to get functional cells (300xg for 5 min at 4°C). Supernatants were separately placed in a fresh tube and centrifuged at maximum speed (13000xg) to obtain a cellular-free *O. vulgaris* serum (OS) as the medium for the next experiments. Cellular pellets were resuspended in a volume of 400 µl of each of the tested media described below and kept at 4°C.

### 2.3. Media and CryoProtectant Agents (CPAs)

To evaluate the effect of different solutions on the cryopreservation process, serum from O. vulgaris (OS), MAS medium, Squid Ringer’s solution (SRS) (Nyholm et al., 2009), and RPMI 1640 (Gibco) medium were used to resuspend the cells. Three biological replicates of each cellular suspension were used to test each candidate media. Two aliquots of each media were used to check the effects of ethylene glycol (EG) and dimethyl sulfoxide (DMSO) as CPAs. For that, one replicate was supplemented with a solution of 15% (final volume) of EG and another with 10% of DMSO. The remained replicate was kept without any CPA and acted as negative control. All aliquots were placed in a Mr. Frosty^TM^ freezing container previously cooled to 4°C and filled with isopropanol, and immediately placed at −80°C to obtain a slow and progressive freezing rate (Mr. Frosty produces a temperature decrease of approximately 1°C min^-1^). This experiment was repeated three times.

### 2.4. Thawing conditions

To determine the optimal thawing protocol, two different methodologies (“slow or progressive” and “fast or non-progressive” conditions) were tested. For the “slow protocol”, cells were thawed in a water bath at 30°C. As soon as the sample showed the first signs of thawing, the original medium without CPA in which the cells were originally frozen was used to slowly dilute the cell suspension by alternating the addition of medium with gentle mixing movements. Cells were slowly transferred into a 50 mL tube (one drop every 5 seconds) and the medium was slowly and progressively diluted up to tenfold. The cells were washed twice by centrifugation at 300xg for 5 min at 4°C. The final pellet was resuspended in 400 µl of each tested CPA-free medium. For the “fast protocol” frozen vials were also placed in a water bath at 30°C and the cells were diluted rapidly and only to twice their volume with CPA-free medium. The cells were pelleted by centrifugation. The supernatant was removed and the cells were resuspended in a final volume of 400 µl of each tested CPA-free medium.

### 2.5. Flow cytometry assays

For quantification of cell viability, a total volume of 100 µL of each thawed cell suspension was incubated in the dark with 2 µL of propidium iodide (PI) (1mg/mL) or 0.5 µL of DRAQ7 (0.3 mM) before measurement by flow cytometry using a Cytoflex (Beckham Coulter), in the Flow Cytometry Service at CINBIO (University of Vigo). PI and DRAQ7 are membrane-impermeable dyes with the ability to stain nuclei that are excluded in viable cells. Their binding to double-stranded DNA emits fluorescence in phycoerythrin (PE) channels (maximum wavelength of 617 nm) for PI or far-red (maximum wavelength of 665 nm) for DRAQ7, which can be detected. PI was used for the initial quantification of the viability after the freezing/thawing of the cells in different media. For the following flow cytometry viability experiments described, DRAQ7 was used instead of IP because of its compatibility to be visualized at the same time in the microscopic analyses. The negative fluorescence threshold and possible autofluorescence emission of hemocytes were previously established and tested using unlabeled cells. Before viability analysis by flow cytometry, cells were also visualized by light microscopy to check their morphology and structure.

### 2.6. Microscopy analysis

To evaluate the morphology of the cells after the freezing/thawing protocol and their viability, different cell suspensions were thawed following the optimal protocol previously established and seeded into an 8-well plate in a final volume of 200 µL of MAS media. To analyze the reduction of the possible stressful effect of the freezing/thawing protocol on the cellular morphology, a cellular batch was thawed 16-18h before the analysis. Cells were seeded in MAS media at kept at 15°C until the image analysis. The morphology of fresh cells, directly thawed cells and thawed and acclimated cells were analyzed under a Thunder fluorescent inverted microscope (Leica). Images were recorded for 4h every 10 minutes and were analyzed to detect differences in the morphology and for the different cellular types.

### 2.6. Phagocytic assays

Once the optimal medium, CPA, and thawing protocol (based on the cellular viability and microscopy morphology results) were selected, cells were frozen and thawed using the optimal conditions. The phagocytic activity of the thawed cells was determined by flow cytometry and by fluorescent microscopy, measuring the ability of the hemocytes to phagocyte a fluorescent and heat-killed inactivated bacteria (*Vibrio lentus*, CECT 5110T, kindly provided by Dr. Rosa Farto from University of Vigo). For cytometry analysis, octopus hemocytes were dispensed into 96-well plates (100 μL per well) and, after 30 min of incubation at 15 °C, a solution of 100 μL of FITC-labelled bacteria with a concentration adjusted to 10:1 (cells:bacteria) proportion was added to the cells. Control cells were incubated with the same volume of filtered FSW. After incubation in the dark at 15 °C for 2 h, the non-internalized particles were removed by washing twice with 100 μL of FSW. The attached cells were finally resuspended in 200 μL of FSW. Twenty microliters of 0.4% trypan blue solution in FSW were added to each sample to quench the fluorescence of adhered but non-phagocyted bacteria. Cells were analyzed in a Cytoflex flow cytometer (Beckham Coulter), in the Flow Cytometry Service at CINBIO (University of Vigo). Phagocytosis was analyzed and it was expressed as the percentage of cells that has internalized fluorescent bacteria (positive cells) and as the median of the fluorescence intensity, which indicated the quantity of fluorescence bacteria phagocyted by each cell. Fresh cells were used as the positive control. To analyze the reduction of the possible stressful effect of the freezing/thawing protocol on the cellular response to fluorescent bacteria, a set of cells was acclimated to the media after thawing for 16-18h by seeding into a 96-well plate and kept at 15°C in the dark before the evaluation of their phagocytic activity.

The phagocytic activity was also analyzed under the Thunder fluorescence inverted microscopy. As for the cytometry analyses, a set of cells was also thawed and acclimated before the experiments. A total volume of 100-200 µL of fresh thawed and acclimated cell suspensions was seeded into an 8-well plate in the presence or absence of a FITC fluorescent labeled bacteria, which was added to the cells in a proportion of 1:10 (cell: bacteria). Images were recorded for 4h every 10 minutes. Image analysis was performed to detect differences in the functional activity of the different cellular types.

### 2.8. Molecular assays

To quantify the effect of the optimized freezing/thawing protocol on the RNA quality and gene expression efficiency, total RNA from cryopreserved cells was isolated and compared with total RNA directly isolated from frozen samples where cells were kept directly in TRI Reagent® (MRC). RNA purification was performed with the Direct-zol RNA Miniprep Kit (Zymo Research), which included a DNase I treatment following the manufacturer’s instructions. The quality of isolated RNAs was tested using a Bioanalyzer (Agilent). A total of 1000 ng of RNA was used to be reverse transcripted to cDNA using the Maxima First Strand cDNA Synthesis Kit for RT-qPCR (ThermoFisher). Octopus ubiquitin expression was used as housekeeping gene, (García-Fernández et al., 2016) to check de reverse transcription efficiency. Quantitative PCR (qPCR) assays were performed using Quant Studio 3 (Applied Biosystems). A total volume of 25 μL PCR mixture included 12.5 μL of SYBR Green PCR master mix (Applied Biosystems), 0.5 μL of primers pairs 10 μM (Ubiquitin-OV-qPCR F: 5′-AGAAGGTTAAGTTGGCGGTTTTG-3′ and Ubiquitin-OV-qPCR R: 5′-CCAGCTCCACATTCCTCGTT-3′), and 1 μL of a 1:5 dilution of the cDNA. Amplification was carried out at the standard cycling conditions of 95 °C for 10 min, followed by 40 cycles of 95 °C for 15 s and 60 °C for 1 min. Each reaction was conducted in triplicate.

### 2.8. Statistics

Viability results were analyzed using a three-way ANOVA with medium, type of CPA, and thawing method as fixed factors. Before analyses, normality and homogeneity of variances were checked by Shapiro-Wilk and Levene’s tests, respectively. In case the homogeneity of variances is violated, raw or rank-transformed data were considered to run the ANOVAs (Conover, 2012). Results were considered significant at p ≤ 0.05. The univariate test of significance was used to elucidate how much of the total variation in the dependent variable can be attributed to differences between factor levels and to know which of the factors have a stronger effect on the dependent variable. Homogeneous groups were established a posteriori with Tukey’s tests for multiple comparisons. Statistical analysis was done using the STATISTICA v.7.0 software (StatSoft). Data are reported as means + SE.

## 3. RESULTS

### 1. Screening of the optimal medium and conditions for O. vulgaris hemocytes cryopreservation

Different percentages of viability were found for each analyzed condition (Figure 1). The analyzed medium, the thawing methodology, and the presence/absence of CPA were factors statistically significant by themselves and also in combination among them (Table 1). According to statistical analysis, the medium used for the freezing/thawing process proved to be the most important factor for cellular viability. Cells preserved in MAS showed the highest percentage of viable cells even when the freezing/thawing process was “fast”, presenting higher values of cellular viability than other media used in “slow” conditions. Cells kept on RPMI medium showed the lowest viability values independently of the freezing/thawing method or the CPA used in the experiment. The presence of a CPA in the freezing medium was also crucial for the cells, being the second factor with more statistical weight on the dependent variable. For most of the media, the values of cellular viability when the CPA was present in the medium were higher than in its absence, except in the case of cells maintained in RPMI medium, where the effect of the medium itself was so extremely deleterious that CPAs could not counteract it. No statistical differences were found between DMSO and EG as CPAs for each medium and each freezing/thawing condition. The thawing method was the third most statistically significant factor, and statistical differences between “slow” and “fast” thawing were only observed for cells preserved in RPMI and SRS media.

**Figure 1.**
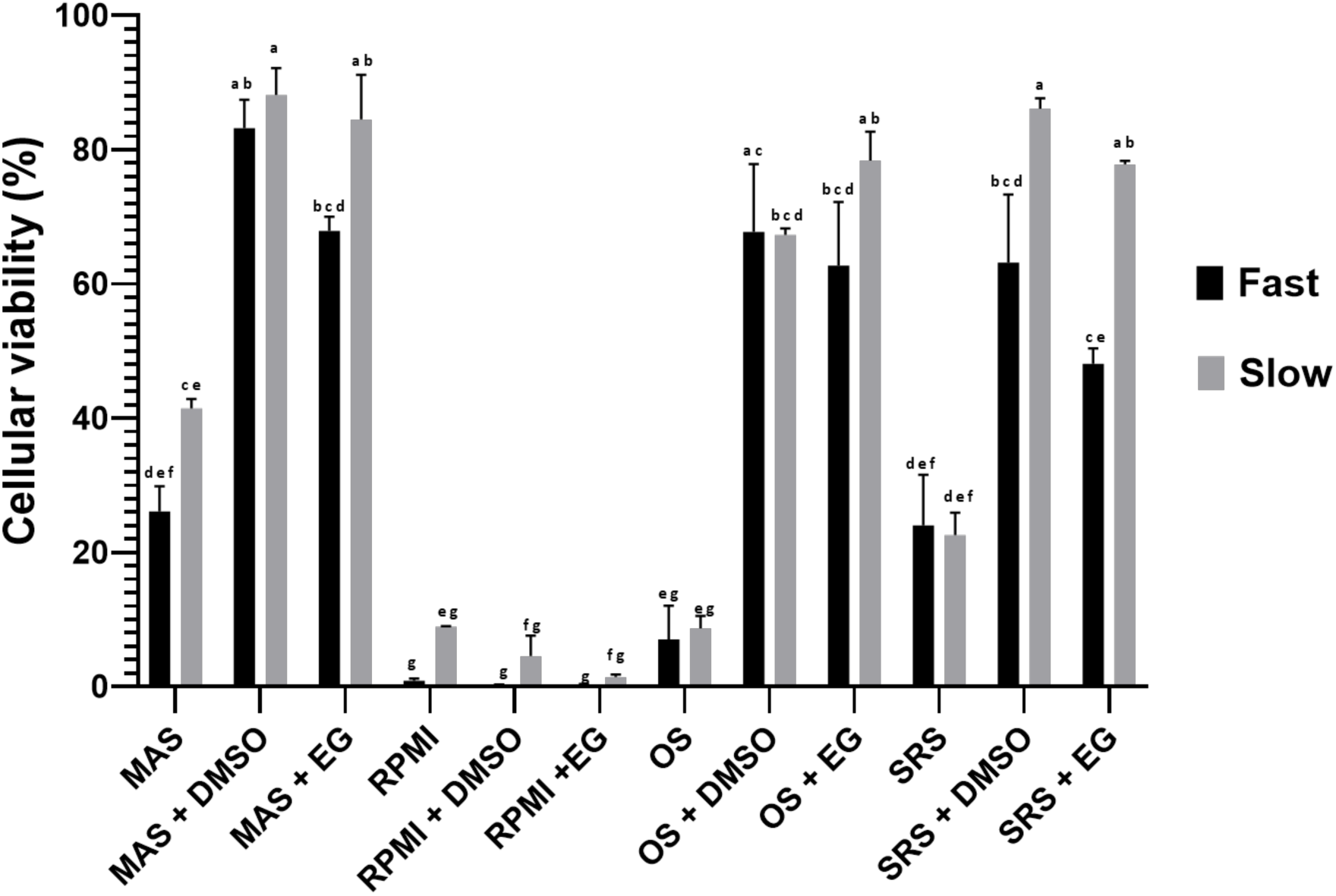
Percentage of cell viability after cryopreservation with the different tested media and thawing following the two evaluated methods. Data represent the mean of values obtained in each experiment + SEM. Bars with different letters indicate significant differences between groups (p ≤ 0.05).

**Table 1.**
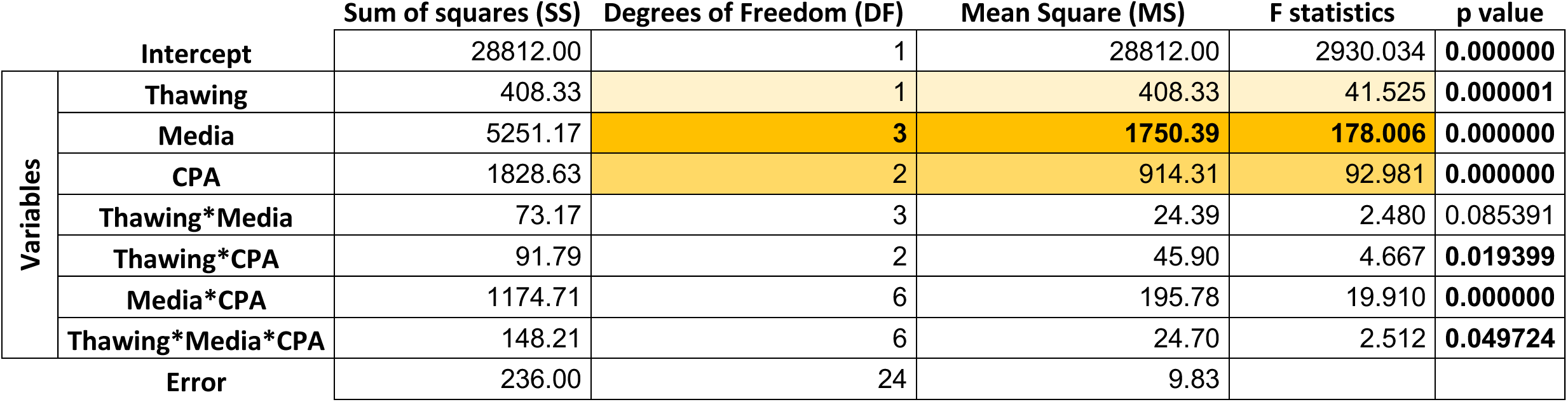
Summary of the univariate multivariate significance tests used in the multifactorial ANOVA.

According to these results, the selected medium for the next experiments was the MAS medium supplemented with CPA (we selected EG instead of DMSO as it is discussed later) and we also chose the “slow” thawing conditions such as is also explained in the next section.

### 2. Cellular survival over time and morphological aspect

Several cellular aliquots were maintained at −80°C, over different periods of time using MAS medium supplemented with EG. After performing the freezing/thawing process following the optimized protocol, the cellular viability was measured by flow cytometry. The viability values were slightly reduced over time, but after 4 months at −80°C, cells still showed viability percentages of more than 75% (Figure 2) suggesting that the freezing and thawing protocol were optimal for this cellular type. The visualization of the cells under the microscope showed that the thawed cells took more time to produce pseudopodia in a similar way to fresh cells. However, after a period of acclimation of 16h, the morphological aspect of the thawed cells was more similar to the aspect of fresh cells, showing an appearance of cellular membrane less rigid than those observed directly after the thawing process. This improvement of the aspect was also observed in the cellular shape, where the acclimated cells did not show a morphology so circular than non-acclimated and their membrane began to produce some movements and invaginations (Figure 3).

**Figure 2.**
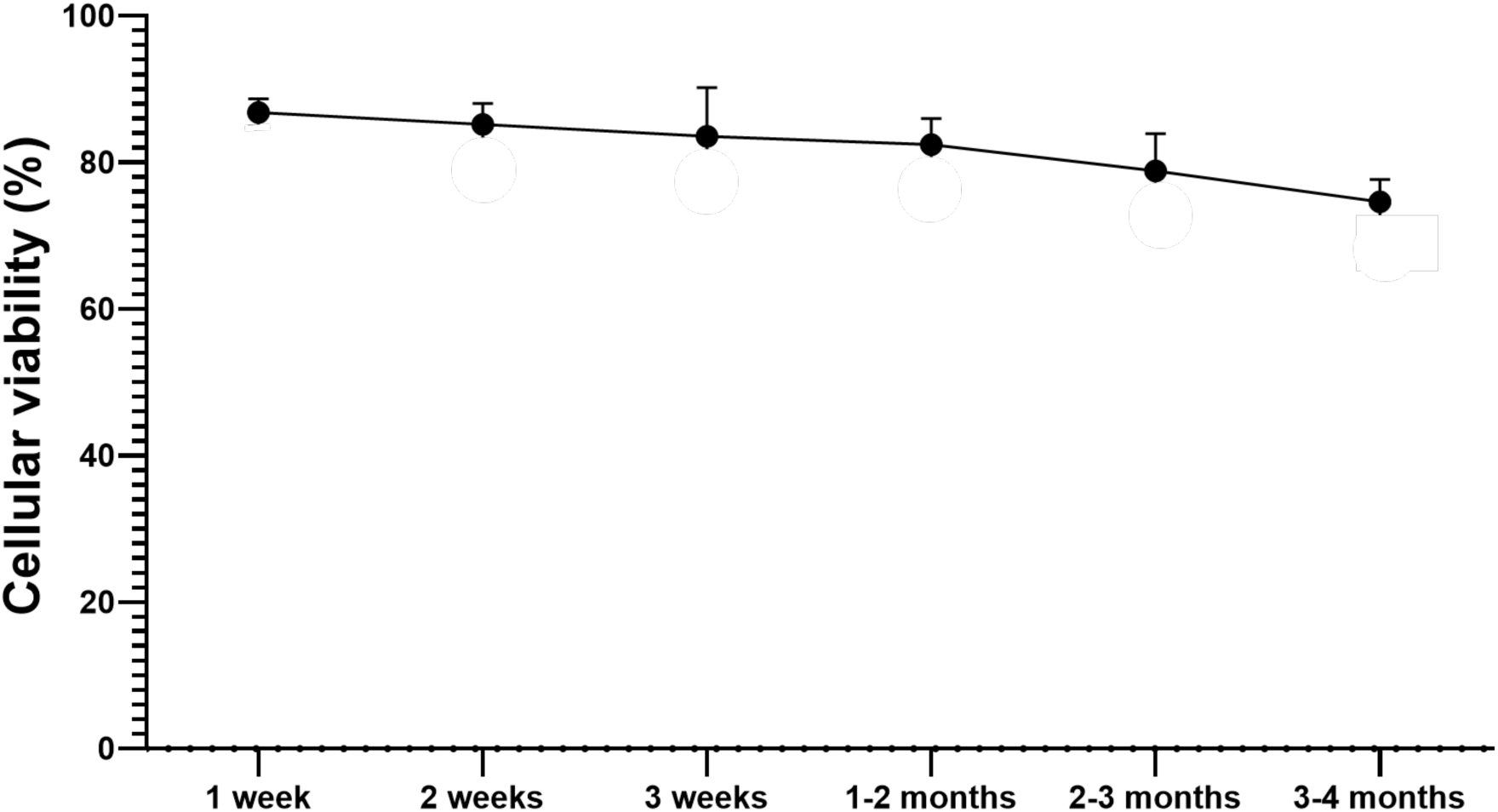
Percentage of cellular survival over time. Data represent the mean of values obtained in each experiment + SEM.

**Figure 3.**
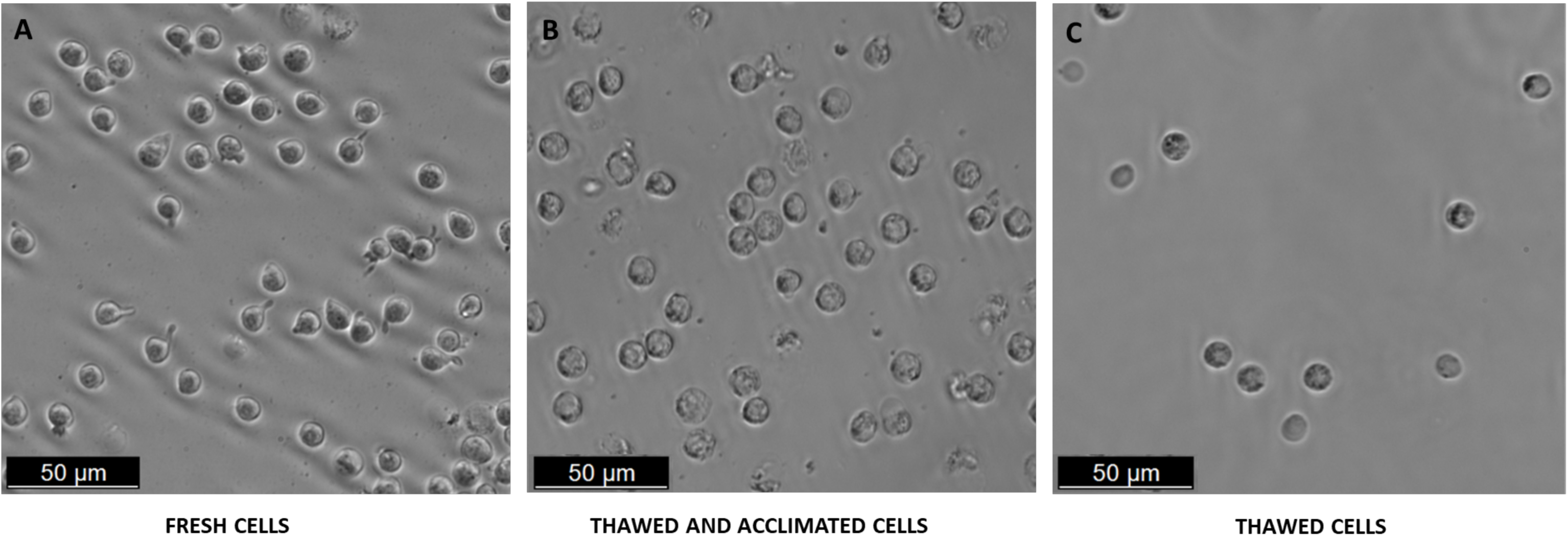
Morphological aspects of fresh (A), thawed and acclimated (B), and directly thawed hemocytes visualized under the optical microscope.

### 3. Functional assays

The functional activity of the thawed cells was measured evaluating by assessing their ability to phagocyte bacteria. The percentage of cells with phagocytic capacity was 100% for fresh cells and close to 90% for the acclimated cells after the thawing process. The phagocytic activity measured immediately in thawed cells (without a period of acclimation) was significantly lower than in fresh cells but did not shown statistical differences with regard to the thawed and acclimated cells (Figure 4A). Despite a similar percentage of phagocytic cells were found between directly thawed cells and acclimated cells, the median of intensity fluorescence (grey bars), i.e., the quantity of phagocyted bacteria by each cell, showed significant differences between acclimated and non-acclimated thawed cells. The fluorescence peaks between cells in the presence (red lines) and in the absence (black lines) of fluorescence bacteria were clearly separated showing that fluorescence events corresponded to cells that had phagocyted labeled bacteria (Figure 4B). However, the intensity of the fluorescence peaks showed variations among the three experimental groups. The maximum peak of fluorescence for fresh cells was located around values of 10^5^ of MFI (median of the fluorescence intensity) whereas the maximum values of the peak for thawed and non-acclimated cells were located in values of 10^4^ of MFI. Curiously, acclimated cells after the thawing process showed a double peak: one close to the maximum values of fresh cells (10^5^) and the other closer to the maximum peak detected for thawed and non-acclimated cells (10^4^), suggesting that the acclimated cells tend to recover their functional activity when they are subjected to an acclimation process after thawing.

**Figure 4.**
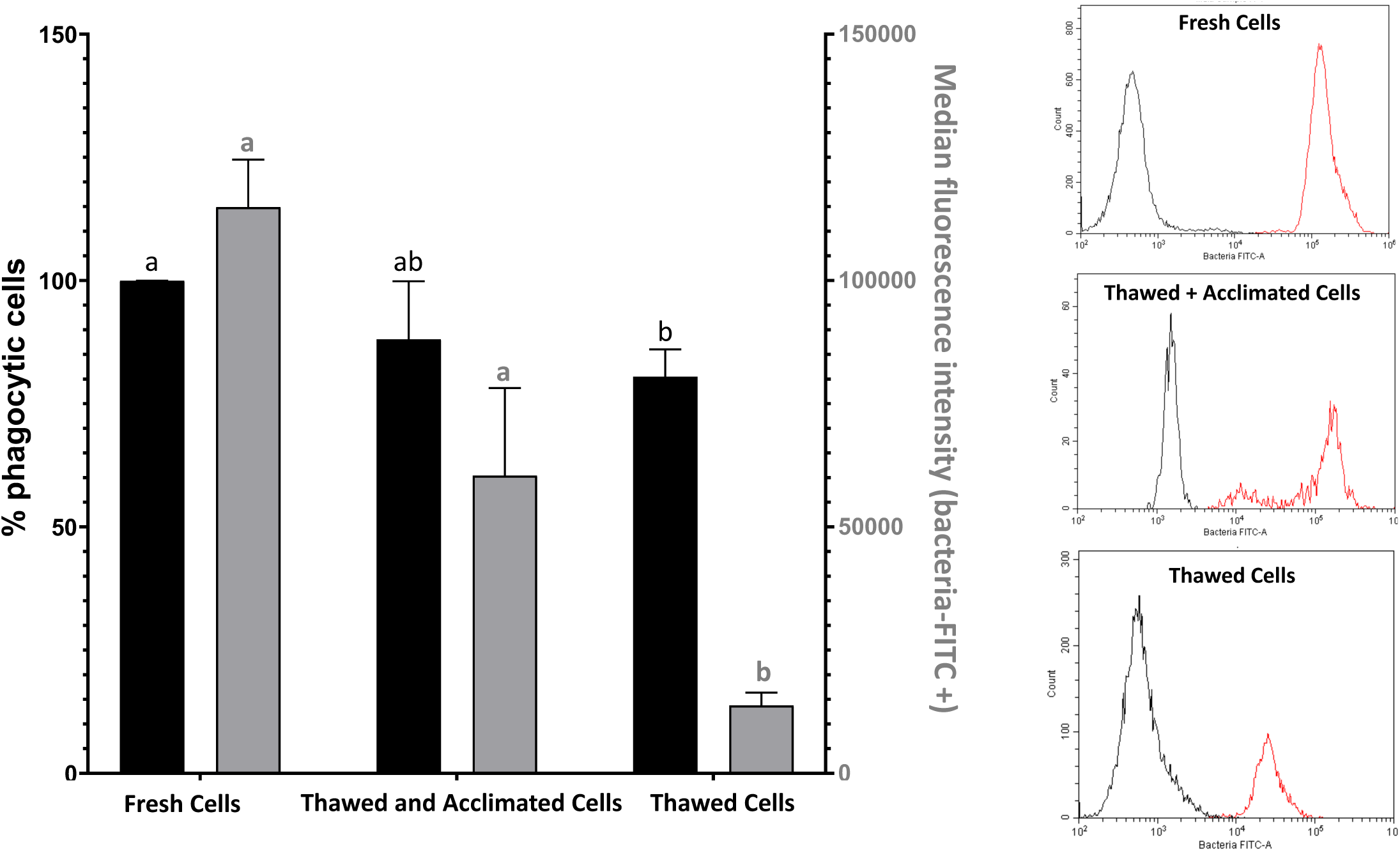
Assessment of cell functionality by measuring cell phagocytic capacity by flow cytometry. Representation of the percentage of phagocytic cells (black bars and “y” left axis) and median fluorescence intensity (grey bars and “y” right axis) of the positive fluorescence population (bacteria-FITC + population) (A). Data represent the mean of values + SEM. Fluorescence cytometry overlay showing the distribution of the total cell population for fresh, thawed and acclimated, and directly thawed cells (B). The red curves represent positive fluorescence, while the black curves indicate negative fluorescence. The peaks of both populations are clearly separated, indicating the efficacy of the fluorescent marker in identifying the subpopulation of interest.

To complement the data obtained in the flow cytometer analysis with a microscopic observation, the results of the visualization of fresh, directly thawed, and thawed and acclimated cells in contact with FITC-labeled bacteria are presented in Figure 5. Both groups of thawed cells showed fluorescent bacteria into the cells although the cells subjected to an acclimation process before the experiment resulted more efficient in internalization of the labeled bacteria, suggesting, once again, the importance of an acclimation process in the recovery of the functional cellular activities.

**Figure 5.**
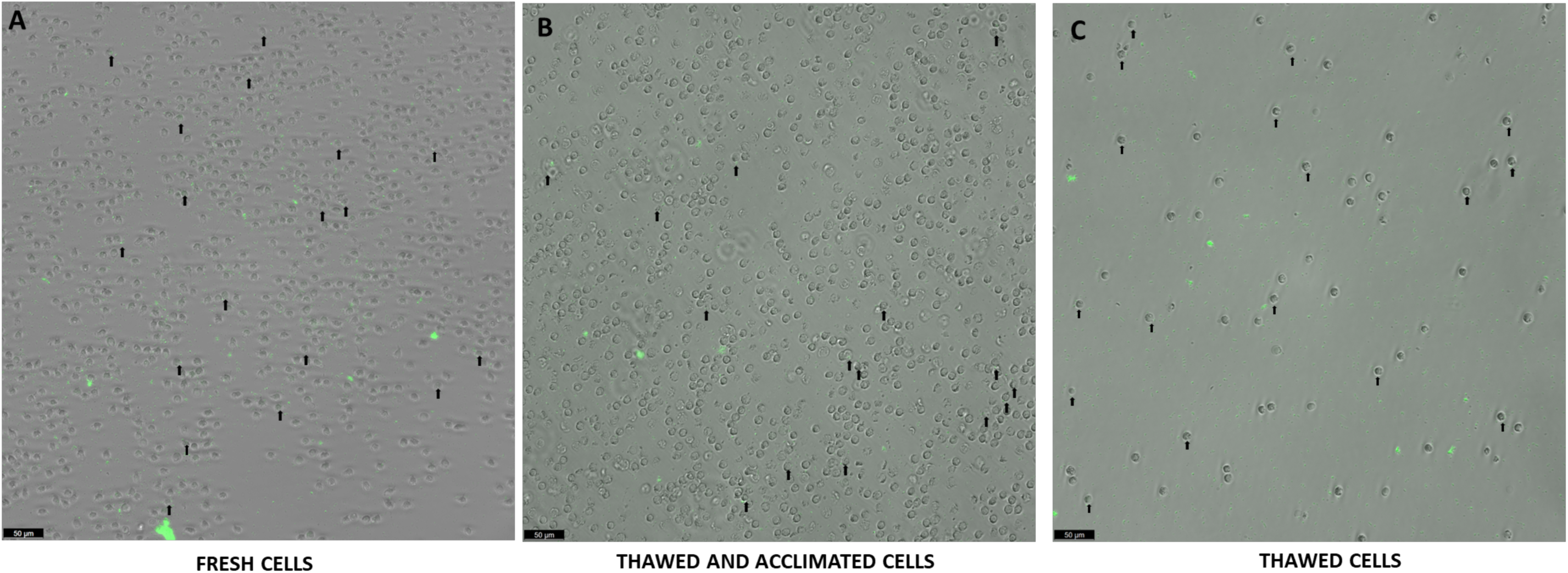
Visualization of the phagocytic activity of fresh (A), thawed and acclimated (B), and directly thawed cells (C) on FITC labeled bacteria. Arrows indicate a representative number of bacteria (green points) located inside the hemocytes.

### 4. Cellular viability

The cellular viability was measured by flow cytometry in four experimental groups: fresh cells (positive control of viability), thawed and acclimated cells, directly thawed cells, and dead cells (negative control of viability) (Figure 6A). The experimental results showed that fresh cells presented values close to 98% and dead cells by osmotic shock showed values lower than 7% of viability. Directly thawed cells presented values close to 75% of viability whereas cells subjected to an acclimation process before thawing showed higher values (close to 85%). A double population of cells was observed in the dot-plot for non-acclimated cells indicating the switch that cells suffer from the live-to-death status and suggesting, the importance of acclimation for the maintenance of viability (Figure 6B). A visual screening was also performed under the microscope using a death-cell marker (Figure 7). The images confirmed the previous results observed by flow cytometry.

**Figure 6.**
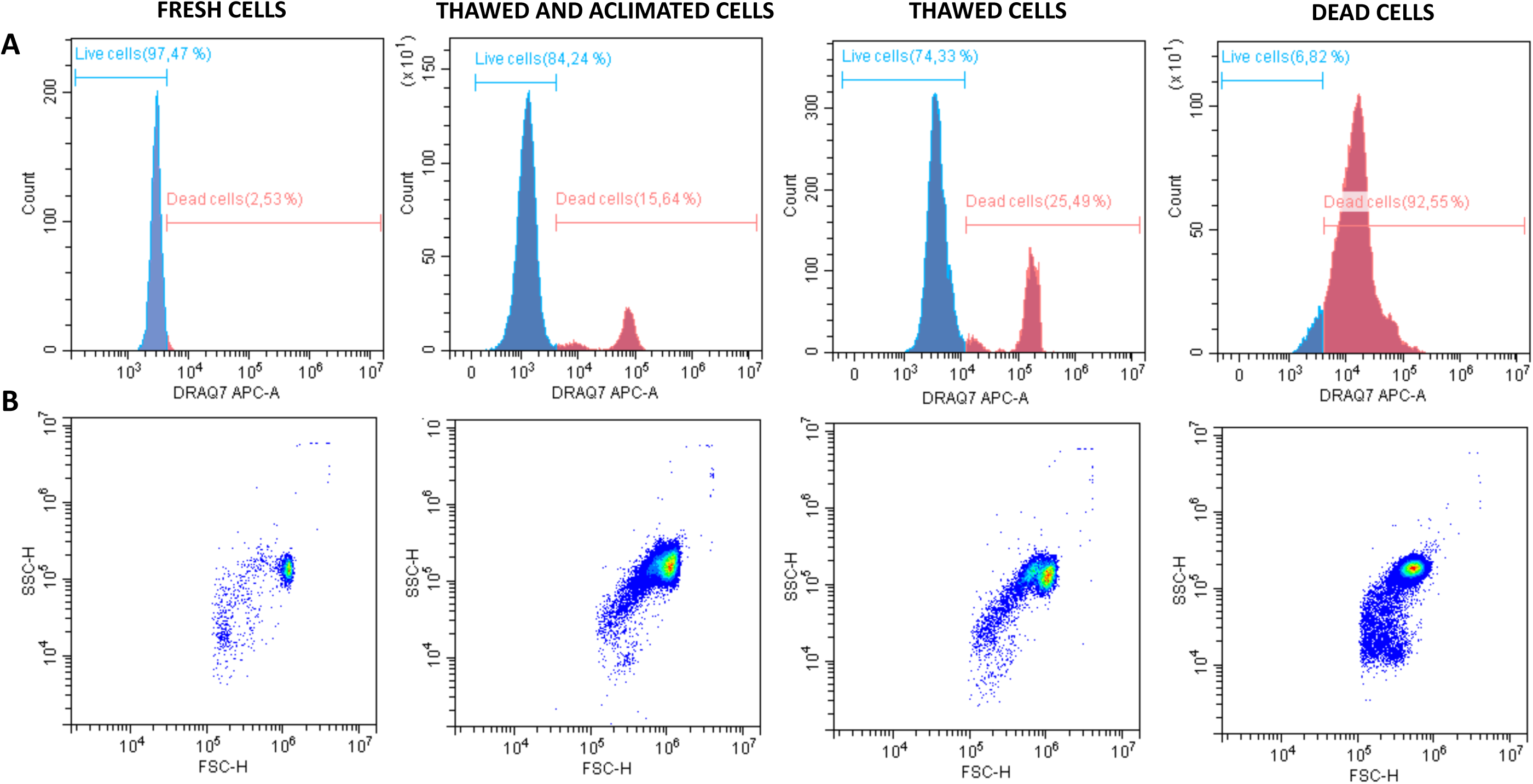
Histograms and dot plots of fresh, thawed and acclimated, directly thawed, and dead cells previously labeled with DRAQ-7 as death marker fluorophore. Histograms represent the distribution of the counts on the fluorescence range. Blue curves correspond with live cells whereas red curves indicate the dead cells (panel A). Dot plots showing side scatter (SSC) *vs* forward scatter (FSC) for the four experimental analyzed groups (panel B).

**Figure 7.**
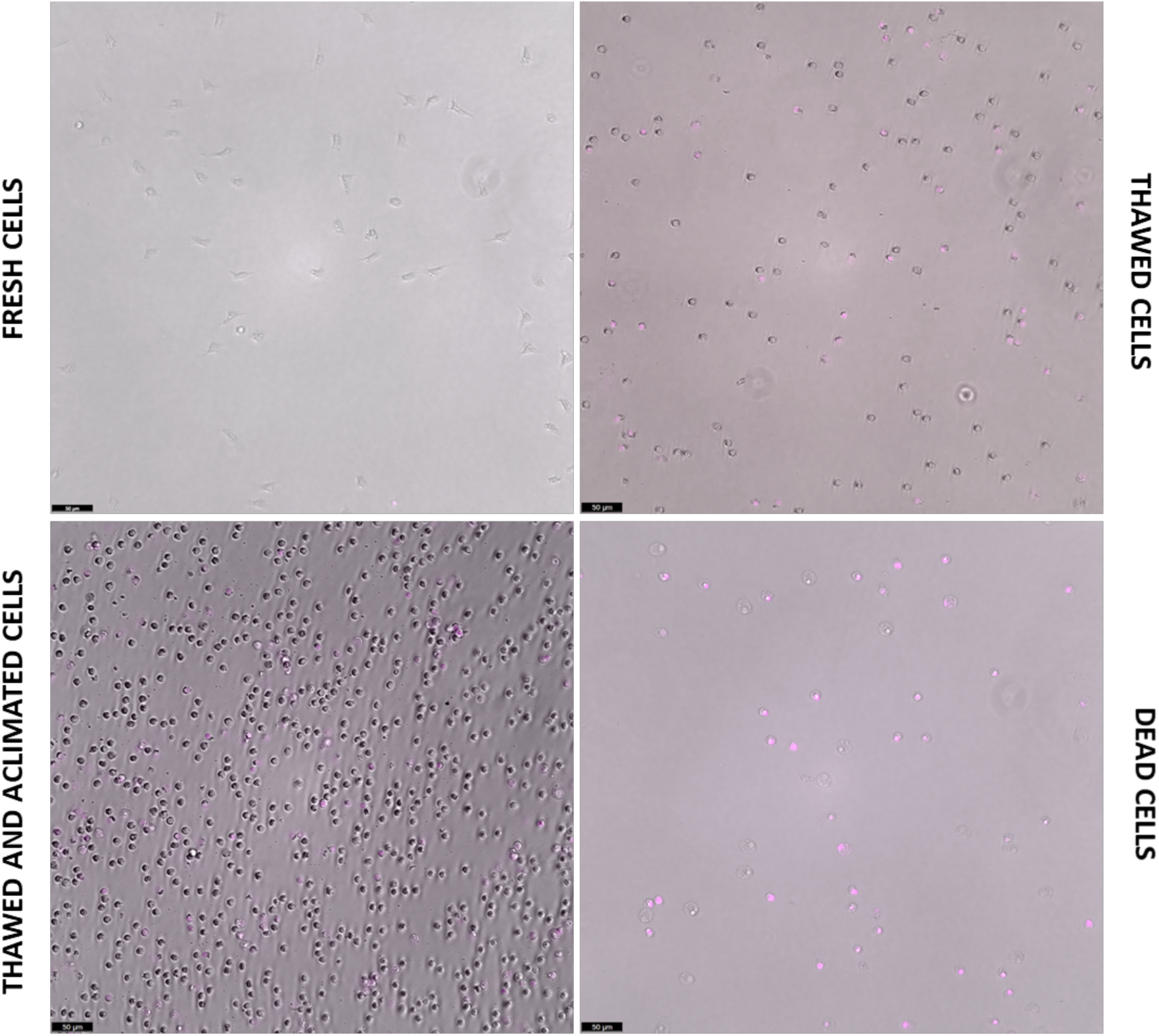
Visualization under the optical microscope of live, thawed and acclimated, directly thawed, and dead cells previously incubated with DRAQ-7 as death marker fluorophore. Pink points correspond with death-positive cells.

### 5. Molecular assays

The RNA integrity profile (RIN) for samples of cells preserved in MAS medium supplemented with EG and for cells preserved in a known safety RNA medium such as TRI Reagent® was analyzed (Figure 8A). Both graphics showed a similar pattern and the expected peaks corresponding to 18 and 28S RNAs were clearly differentiated. No signs of degradation were detected for any sample. To detect any type of interference of MAS medium on the gene expression detection (at RNA isolation, reverse transcription, or qPCR levels) the ubiquitin was amplified by qPCR as a housekeeping gene, (Figure 8B). The amplification curves were very similar, and even overlapped completely from cycle 20 onwards, showing that the MAS medium does not seem to interfere in the gene expression.

**Figure 8.**
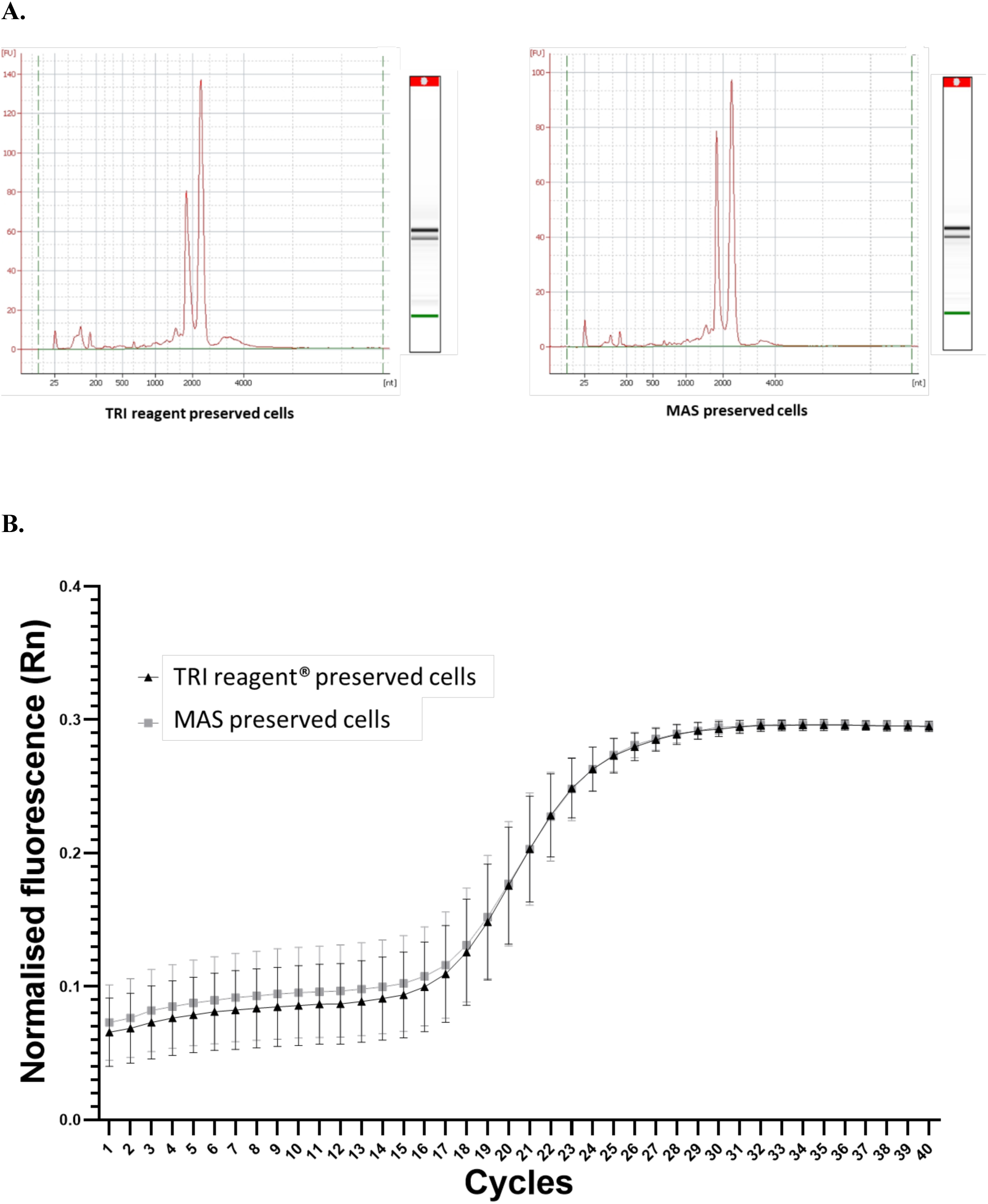
Profile obtained from an Agilent Bioanalyzer. The x-axis represents the migration time (seconds), while the y-axis shows the fluorescence intensity, indicating the concentration of RNA present for samples cryopreserved in TRI reagent and in MAS medium (A). Amplification curve from qPCR analysis of the housekeeping gene, ubiquitin (B). The graph shows the cycle number on the x-axis and the fluorescence intensity on the y-axis. The triangles line represents the amplification curve for TRI reagent® preserved cells, whereas the squares line indicates the amplification for MAS cryopreserved cells.

## 4. DISCUSSION

In the cryopreservation field, the selection of the optimal medium for a specific cellular type, as well as, the choice of the appropriate CPA and the methodology used for the freezing/thawing steps (cooling and warming rates and temperatures) is specific for each cellular type. The media and employed methodologies may be ideal for one type of cell while being completely deleterious for another. For instance, a CPA as glycerol is widely used for cryopreservation of sperm from different species (Sieme et al., 2015) and also for human red blood cells (Rogers et al., 2018). However, many types of cells are difficult to be cryopreserved using glycerol as CPA. The reason why some cell types one protectant works better than others seems to be related to cell-specific cytotoxicity and membrane permeability and other unclear factors. Therein lies the difficulty of finding specific protocols that manage to maintain the functional and molecular characteristics of the cells under study. A complete cryopreservation formulation should consist of several components, such as the presence of mineral salts, buffering components, specific nutrients for the cell type to be studied, CPA agents, antioxidants, and scavengers (Wolkers and Oldenhonf; 2015). In the present paper, we have tested a total of 4 media without or in the presence of two different CPAs: EG or DMSO. Our results revealed that the MAS medium reported the best viability results after freezing and thawing of the cells, showing values close to 90% when it was supplemented with a CPA. Among both tested CPAs, no significant differences between DMSO and EG were found. Despite DMSO has been widely used as CPA for many animal and vegetable cells, some studies have revealed its possible role as cellular apoptosis and stress enhancer (Kang et al., 2017; Blanco et al., submitted for publication). In addition, EG has been reported as a proper CPA for other marine organisms, molluscs in particular (Heres et al., 2019; 2021; 2022), with good rates of cell and larvae survival, as well as to avoid the presence of larvae malformations. Although the cryopreservation medium must be optimized for each biological material in a specific way, the lack of statistical differences found between the two tested CPAs and the results documented in the literature tipped the balance towards the use of EG for octopus hemocytes cryopreservation. However, it is worth to note that, for this specific case and, based on the viability results obtained, either of the two could have been used as CPA. The MAS medium itself (without any CPA) was able to produce better viability values than other tested media even supplemented with CPAs, suggesting that the MAS composition counts with essential components for the conservation of the cells. Concretely, the MAS medium contains glucose, which provides the energy for basic maintenance and cell metabolism during the cryopreservation period and, as happened with other sugars, provides antioxidant protection (Pan et al., 2023; Whaley et al., 2021; Anjos et al., 2021); trisodium citrate, which supplements the medium with mineral salts providing a suitable ionic environment that mimics the natural extracellular environment, maintaining the osmolarity and ionic balance of the cells while acting as a buffer, keeping the pH stable during the freezing and thawing process and thus protecting the cells from sudden changes in acidity or alkalinity that could damage them (Herrler et al., 2006). Moreover, this molecule also can act as a scavenger by binding to metal ions such as calcium and magnesium, which helps to prevent the formation of precipitates and stabilize solutions (Pham and Schwartz; 2013). In the specific context of hemocytes, as part of their defense mechanisms tend to aggregate to catch microorganisms and fight more efficiently against them (Gliński and Jarosz; 1997). These aggregates, although advantageous for the pathogens fighting, pose a problem in cryopreservation since the cells do not receive the cryoprotective medium equally. Also after thawing, cells need to be individualized for subsequent analyses, including single-cell research. MAS medium also contains citric acid, which plays a multifaceted role in cryopreservation, helping to maintain pH, sequestering metal ions, reducing oxidative stress, improving cell permeability, and stabilizing biomolecules (Fuller, 2004; Patel et al., 2016). The MAS formula is completed with EDTA and sodium chloride, which proportionate a chelating agent, EDTA (Masoudi et al., 2020) and the proper salts, sodium chloride, for keeping alive marine cells. No significant differences in cellular viability were neither found concerning SRS medium (Nyholm et al., 2009), which also contains mineral salts and chelating agents in its composition. However, our microscopic observations before and after the freezing/thawing process (data not shown) revealed more debris and a worse cellular aspect of the cells resuspended in this medium. Good viability results were also found when the cells were resuspended in OS. Nevertheless, it is worth to note, that for obtaining the octopus serum, the hemolymph extraction was conducted in a proportion 1:1 with MAS medium as anti-aggregant solution to avoid cellular clotting, thus the OS was diluted in MAS medium proportionating probably the salts, chelating elements, and other optimal molecules, which would facilitate the cryopreservation process, although the viability values were significantly lower in the case of the use of DMSO as CPA. Other different culture media were also tested, although cell viability found was extremely low (data not shown). Surprisingly, L-15 medium, which has been referenced in multiple papers on primary cultures of marine cells (Costa et al., 2012; 2013) produced very high mortality in our cells. This medium was also used by Songco-Casey et al., (2022) in their studies of *Octopus bimaculoides* visual system and by Styfhals, et al. (2022) in their analysis of the cellular diversity of *O.vulgaris* brain. However, in the case of *O. vulgaris* hemocytes, the results were not as satisfactory. This shows, once again, that each cell type needs to be preserved in a specific way. For all these reasons our selected medium was MAS supplemented with EG.

In addition to the CPAs, cryopreservation also requires optimal cooling and warming rates to control cellular dehydration during freezing, the intracellular ice formation, and also the possible deleterious effect of CPAs during the thawing process (Wolkers and Oldenhonf; 2015). For the last point, according to previous data obtained for marine molluscs, we have used a cooling rate of −1°C/min (Heres et al., 2022) and it is also very important a progressive dilution of the freezing medium to dilute as much as possible the CPA and prevent the cells to damage. Different researches have shown that progressive dilution of CPA during the thawing process is superior to rapid thawing for maintaining cell viability allowing a gradual equilibration of osmolarity, reducing the risk of osmotic shock, which is detrimental to cell membranes and overall cell survival (Baust et al., 2017; Agarwal and Singh; 2023). Progressive dilution avoids sudden osmotic changes that can occur during rapid thawing, which improves the recovery and viability of cryopreserved cells. Uhrig et al., (2022) collected in their review that optimized thawing protocols, including progressive dilution of CPA, significantly improve the post-thaw recovery and viability of induced pluripotent stem cells (iPSCs). In our case, we tested the fast (non-progressive) and slow (progressive dilution) thawing of the cells for the four tested media. Despite the results previously reported in the bibliography evidenced the lethal effects of a fast addition of CPA-free medium to a recent thawed cellular suspension, that effect was not significantly observed for those cells kept in MAS or OS media, suggesting that the protective effect of the medium, was able to compensate the deleterious effect of the fast process itself, reinforcing the advantages of this medium on the preservation of the cellular viability. For the other tested media, fast or slow processes produced significant differences highlighting the importance of the used methodology. However, in the present case, the MAS medium seems to be potent enough as a cryopreserving medium.

Although cell viability showed a progressive reduction over time, the MAS medium was able to maintain percentages of viable cells above 70% after 3-4 months of freezing at −80°C. It has been reported that cell freezing time can affect cell viability. Several studies have investigated how the duration of frozen storage influences cell viability. Baust et al., (2017) showed that the viability of CD34+ stem cells decreases with freezing time. After a prolonged period of storage in liquid nitrogen, a significant increase in early apoptosis was observed, which negatively affects hematological recovery after transplantation. Li et al; (2021) evaluated the impact of cryopreservation on peripheral blood mononuclear cells (PBMC) from pigs and found that, although the overall viability of frozen and thawed cells remained high, there was a significant decrease in cellular functionality, such as the ability to proliferate after antigenic stimulation. Our data agree with the reported works but it is important to note that, in our case, cells were cryopreserved and then maintained at −80°C, where temperature is not too low to minimize biochemical reactions that can damage cells during prolonged storage but where there is a risk of ice crystal formation that can perforate cell membranes and damage intracellular structures along time. Temperatures of −196°C, molecular movement is extremely low, which reduces cellular damage. Temperature fluctuations are minimal at −196°C, and chemical reactions and degradation processes are completely slowed down, which helps to maintain the integrity of biomolecules in stored cells, and better preserve physiological functions (Baust et al., 2017; Li et al., 2021).

Because the main objective of this work was to provide a greater margin of time to carry out laboratory tasks that cannot be carried out in the same day or even those that must be carried out between different laboratories and that require the shipment of samples, we have only used freezing and storing at −80°C, the results obtained seem to be very promising, suggesting that, at lower temperatures, the results could be even better. Nonetheless further experiments will be needed to figure out the survival rates of the cells after to be kept in liquid nitrogen.

One of the key points of this work was to be able to preserve the cells, not only in a viable state but also in a functionally active state. Considering that preservation at −80°C is not as long-term stable as in liquid nitrogen, the next aspect to be taken into account was to evaluate their functional capacity after a given time. Results obtained presented a good quality of the cells stored for as long as three-four months, presenting a 75% viability. We wanted to compare not only the functionality of the cells immediately after thawing but also their capacity to phagocyte when they were acclimated for some hours to the new conditions. The ability to phagocyte is one of the tools used by the defense cells of these animals to fight against pathogens, as has been previously reported (Castellanos-Martinez et al., 2014b). Our results evidenced that those cells subjected to an acclimation process presented both a better aspect of their membrane showing higher flexibility (visualized under the optical microscope) and also a higher ability to phagocyte fluorescence particles. These results agree with other previously reported data where has been already shown that acclimatizing cells after thawing can improve their functionality. Kuerten et al., (2012) focused on cryopreserved PBMCs and demonstrated that a short pre-culture or “resting” period after thawing can improve the cells’ immunogenicity, allowing the cells to recover from the stress of the freezing/thawing process and regain functionality more effectively. Ortiz-Rodriguez et al., (2019) demonstrated that the functionality of thawed stallion spermatozoa improved significantly through post-thaw incubation with a specific medium. Bellas and Paredes, (2011) after cryopreserving sea urchin embryos observed a period of 24/48 hours of delayed development immediately after thawing, which was presumably used to recover from stress and repair minor damages, followed by a normal larval development. This treatment activated pro-survival pathways and enhanced mitochondrial function, reducing cellular stress and apoptosis, which are critical for maintaining cell functionality. These findings suggest that a period of acclimation after thawing can be beneficial, helping cells to recover and perform better in subsequent applications. In our case, we observed that cells subjected to acclimation were able to present better values of phagocytosis, both in the percentage of phagocytic cells and in the median of the fluorescence intensity, i.e., more percentage of cells were able to phagocyte and more bacteria were phagocyted per cell when the thawed cells were previously acclimated to the new conditions. Our results also showed visual evidence of this recovery where, on the one hand, we observed that cells subjected to post-thaw acclimatization show a fluorescence peak at the same height as that observed in fresh cells and another similar to that observed in thawed and non-acclimatized cells, suggesting that they are recovering their phagocytic capacities during this period of acclimation. On the other hand, photographs taken under the microscope showed that acclimated cells had a greater flexibility of their membrane, which translated into a greater phagocytic capacity, as well as, higher levels of viability. In addition, flow cytometry analysis revealed the presence of a double population (dead and alive cells) in the case of non-acclimated cells, suggesting that this “resting” period can contribute to increasing cellular viability. All these results are evidence of what has been described in other species and demonstrate that cell acclimation after thawing is key to recovering its membrane, functionality, and viability.

In addition to ensuring that cryopreserved cells are viable and functional, another important objective of this work was to maintain live cells with quality for carrying out molecular assays such as scRNA-seq analysis. As we have already mentioned previously, this methodology requires being able to start from a population with a high percentage of viability and where the cells are individualized and not grouped. These requirements seem to be achieved after the results previously discussed, but for a gene expression analysis, it is essential to be able to start from a quality RNA, which is intact and where there is no degradation or contamination (Haque et al., 2017). To solve this last point, we wanted to analyze the RNA integrity of cells cryopreserved in MAS medium in comparison with TRI Reagent®, a medium that has been proven effective in maintaining RNA integrity during storage, without significant adverse effects on gene expression (Ma et al., 2010). The lack of differences between both RIN patterns demonstrates that MAS medium is also available as an RNA integrity preserver. The quality of the population of total RNA can also be evaluated through the measurement of the expression of a housekeeping gene after reverse transcription-quantitative real-time polymerase chain reaction (RT-qPCR) (Caballero-Solares et al., 2022). The efficiency of reverse transcription depends on the integrity of the initial RNA molecules and the final results in the qPCR. The Ct values obtained (number of cycles where the fluorescence obtained in the analyzed samples is higher than the established threshold) will be lower when the higher the expression of that specific gene is obtained; that is: the higher the integrity of the starting RNA, the lower they will be. Our results revealed that the Ct values for the housekeeping expression analyzed were similar for both tested media, suggesting that MAS media can preserve the quality and integrity as an optimal preserver as TRI Reagent® for common octopus hemocytes.

In conclusion, the data presented in the current paper show a new methodology for cryopreservation of the main immune system cells of *O. vulgaris*, an important model species for a multitude of biological studies and an excellent candidate for aquaculture diversification. The reported data demonstrate the viability, functionality, and molecular integrity of cryopreserved octopus cells, making these cells useful for multiple assays. The successful cryopreservation of these cells represents a powerful new methodology that will allow the development of techniques impossible to perform so far and will facilitate the study of the defense mechanisms of a highly evolved species strategically placed in evolution. Moreover, some of the most cutting-edge molecular techniques, such as scRNA-seq, do not yet offer standardized protocols for the optimal preparation of marine cells. In this work, we propose the possibility of using cryopreservation as a way to keep cells viable and functional while maintaining molecular integrity for this type of assays. Given the lack of cell lines in marine invertebrates, cryopreservation aids *in vitro* cellular assays and significantly reduces reliance on animal experimentation. This not only decreases the number of animals required for research but also aligns with ethical guidelines aimed at reducing animal use in scientific studies.

## ACKNOWLEDGEMENTS

This work was supported by Project PID2020-119906GB-I00 (INMUNOCTOPUS: Immunity in common octopus: non-self recognition and immune response induced by pathogens”) funding by MINECO-Proyecto de I+D+I “Generación del conocimiento” 2020. Estefania Paredes holds a Juan de la Cierva Incorporacion grant funded by MCIN/AEI/10.13039/501100011033 and by “European Union NextGenerationEU/PRTR”. Authors thank to the SACUIM and Microscopy Service of the IIM-CSIC for their technical support.

